# Gait adaptation to asymmetric foot-ground compliance applied by robotic footwear

**DOI:** 10.1101/2025.09.19.677364

**Authors:** Mark Price, Wouter Hoogkamer, Meghan E. Huber

## Abstract

Perturbing foot-ground interaction dynamics has shown promise for eliciting adaptations to inter-limb weight-bearing symmetry, a critical target for rehabilitation of asymmetrical neuromotor deficits affecting gait. To date, this perturbation paradigm has been delivered via adjustable stiffness treadmills, in which the treadmill deck displaces under the walker’s foot during the stance phase of gait. Recently, we developed robotic footwear capable of delivering asymmetrical ground stiffness perturbations during walking, offering a portable experimental platform for asymmetrical ground stiffness perturbations, allowing it to be executed overground and on conventional treadmills. In this study, we quantified kinetic and spatio-temporal gait adaptation to a foot-ground compliance perturbation delivered by the novel robotic footwear. We found that participants shifted their vertical ground reaction force impulse, vertical pushoff peak, and peak braking force toward the unperturbed limb after the perturbation was removed, indicating a shift in neuromotor control elicited by the perturbation. We conclude that the robotic footwear can elicit weight bearing adaptations similar to adjustable stiffness treadmills, and that the dissipative mechanical properties of the shoes likely play a key role in the direction of the adaptation.

## I. INTRODUCTION

Gait asymmetry resulting from hemiparetic stroke is a major cause of adult disability. 80% of people who survive acute stroke go on to experience gait dysfunction [1], and this dysfunction frequently manifests as persistent motor deficits on one side of the body [2]. Aside from being indicative of impaired neuromotor function, asymmetrical gait is associated with slower gait speed [3], increased fall risk [4], and increased risk of developing osteoarthritis [5], [6] and osteoporosis [7].

A promising technique to restore some of this function is implicit learning through error augmentation training [8], [9]. Split-belt treadmills, on which the belt under the paretic limb runs at a different speed than the belt under the nonparetic limb, exaggerate existing step-length asymmetry during exposure to asymmetric belt speeds. This exaggerated asymmetry elicits a neuromotor adaptation which persists after the perturbation is removed, resulting in more symmetrical walking until the effect washes out in a few minutes of walking [8]. While the effects are at first short-lived, repetitive bouts of splitbelt treadmill training have produced improvements in gait symmetry that persisted for at least 3 months [9]. However, while split-belt treadmill training has shown the ability to improve spatial symmetry in gait, it generally does not improve temporal and kinetic gait symmetry [10], [11].

A newer asymmetric gait adaptation paradigm has emerged in the form of asymmetric stiffness treadmill walking [12]– [14], for which preliminary results indicate that adaptations in gait kinetics can be elicited when ground stiffness is perturbed asymmetrically [13], [15]. In particular, targeted changes in symmetry in the vertical ground reaction force (vGRF) at push-off have been observed in response to 10 minutes of exposure to walking on asymmetric surface stiffness [15]. Despite the promise of this finding, adjustable stiffness treadmills possess inherent limitations; there is evidence that motor adaptations to perturbations are context specific, and do not fully transfer to contexts that are different than the training context (i.e., overground walking out of the lab) [16]. Additionally, while adaptations to push-off vGRF were observed, no changes in propulsion were observed, which is a critical functional measure in hemiparetic stroke [17]. It has been suggested that a possible explanation for this lack of change in propulsion force is the constraint of the perturbation to the vertical axis [15]. Finally, should asymmetric stiffness training prove to be an effective rehabilitation intervention for asymmetric neuromotor gait deficits, the hardware required to deliver the intervention is not widely available and is not portable; four prototypes exist and all weigh in excess of 50 kg [12].

To address some of the inherent limitations of asymmetric stiffness treadmill paradigms, we developed robotic footwear capable of delivering a similar perturbation [18]. This platform is portable and lightweight (600 g per foot, 5 kg backpack), and can change the compliance under each foot independently through pneumatic actuation. This technology enables the capability of perturbing surface stiffness overground, potentially addressing the context specificity limitation of treadmill-based paradigms. It also enables adaptation experiments on a fully instrumented treadmill, which circumvents the unique challenge of three-dimensional ground reaction force sensing on the moving platform of an adjustable stiffness treadmill, which none of the existing prototypes have yet accomplished [12], [13]. However, there are inherent differences between a compliant treadmill and our adjustable compliance shoe sole, primarily that the footwear controls a soft, dissipative nonlinear compliant interface compared to the linear stiffness of existing compliant treadmills, and that the compliant interface being worn on the body necessitates a tall platform sole between the foot and the ground (*>* 5cm) to allow for substantial deflection at low stiffness. Therefore, we sought to quantify gait adaptation to a ground stiffness perturbation applied by the novel robotic footwear.

**Fig. 1.**
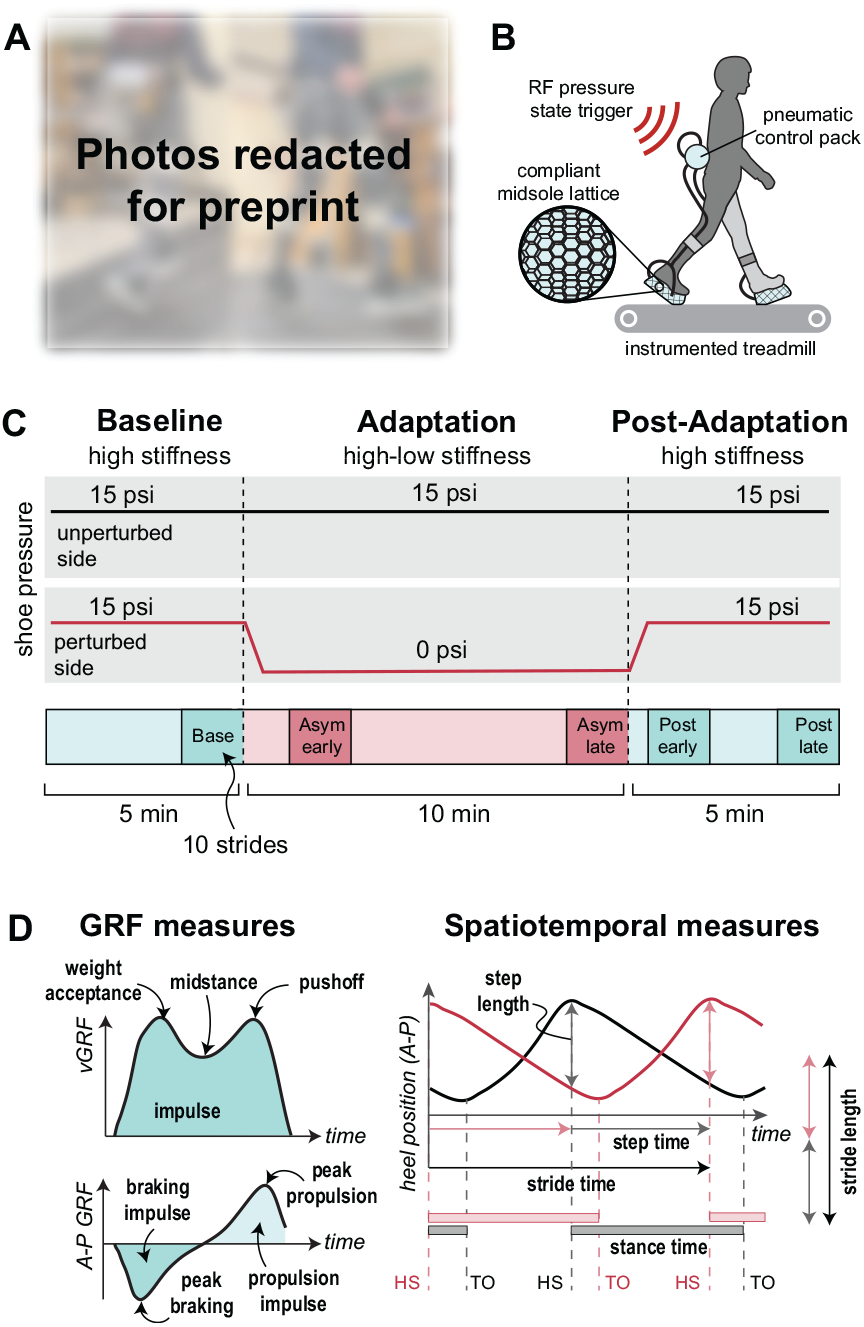
Experimental setup and paradigm. **A:** Photographs of an experimental participant undergoing the experiment. Motion capture markers other than those required for spatio-temporal measurements were not analyzed for this study. **B:** Diagram of experimental setup. Participants walked on an instrumented dual-belt treadmill and experienced asymmetric surface compliance while walking with custom robotic footwear. The footwear are part of an untethered system which control foot-ground compliance inversely with pneumatic pressure, triggered between programmable compliance states with a radio frequency signal (ACFT; [18]). **C:** Experimental protocol. Participants walked for one 20minute bout. The footwear maintained symmetrical, high stiffness (low compliance) for the first five minutes, after which the pneumatic pressure was abruptly lowered on one side for 10 minutes. Symmetrical high pressure was restored for the final five minutes of walking. Effects of the perturbation were assessed in 10-stride windows at the start and end of each compliance phase, with delays to account for the time constant of the compliance change. **D:** Definition of dependent measures. Transient features and impulse were computed for ground reaction forces (GRF). Heel position and heel-strike/toe-off gait event timing were used to compute spatio-temporal measures.

We performed an experiment with neurologically intact young adults to quantify adaptation to asymmetric compliance imposed by the novel footwear. Informed by asymmetric ground stiffness walking experiments on a treadmill [15] and predictive simulations of optimal gait under energetically conservative vs. dissipative surface compliance asymmetry [19], we hypothesized that vertical ground reaction forces and braking/propulsion ground reaction forces would increase on the unperturbed side relative to the perturbed side immediately after the asymmetrical surface compliance perturbation was removed. Our hypothesized asymmetry is in the opposite direction from the treadmill-based experiments, in which the low-stiffness belt had underdamped dynamics. This is because in simulation, ground reaction force measures are larger on the perturbed than the unperturbed side when low stiffness is accompanied by low damping. The asymmetry shifts to favor the unperturbed side in the presence of a high damping coefficient, which more closely resembles the dynamics of the compliant footwear. In addition to ground reaction force measures, we investigated spatio-temporal adaptations to compare adaptations to asymmetric footwear compliance with prevalent adaptations to split-belt treadmill walking.

## II. EXPERIMENTAL METHODS

### A. Participants

A total of 15 participants were recruited for this study. Due to equipment malfunctions in the early collections, 12 participants completed the experiment with all necessary data collected. Participants self reported sex, age, and height, and their mass was measured via treadmill force plates (Sex: 6F/4M/2X, Age: 25 ± 6 years, height: 166 ± 5 cm, weight: 69.2 ± 9.4 kg). All participants had no history of biomechanical, cardiovascular, and/or neurological disorder or injury, were capable of walking at a comfortable pace for 30 consecutive minutes, and had no prior exposure to asymmetric stiffness walking. Each participant signed a consent form before the first session. Approval for these tests was obtained from the Institutional Review Board at UMass Amherst (Protocol #3638).

### B. Adjustable Compliance Footwear

The adjustable compliance footwear technology (ACFT) is our custom wearable system for modulating the foot-ground compliance under each foot independently through pneumatic actuation of 3D-printed pressure vessels embedded into soft midsole lattices [18]. This prototype system allows for online control of the foot-ground interface dynamics, capable of switching between maximum and minimum compliance (∼50–300 kN/m) within about 4 s. In preliminary tests, the maximum compliance condition resulted in about 1.5 cm of vertical deflection at heel-strike for a 64.5 kg participant walking overground [18]. For this study, we perturbed one side by adjusting the compliance under one foot while the other remained in the rigid condition.

### C. Experimental Protocol

Before the experiment, we placed reflective marker balls participants’ heels to quantify step and stride length. Participants then donned the ACFT by first adjusting the pneumatic control system to fit snugly and comfortably to their waist, then strapping the inflatable soles to their existing shoes. Participants were instructed to wear low-profile casual or athletic sneakers they were comfortable walking in to the experiment.

Particpants stood quietly in a neutral posture for 10 s while standing on a force-instrumented dual-belt treadmill (Bertec, Columbus, OH, USA) to obtain their weight while wearing the ACFT. They then performed a continuous 20-minute walking trial at 1.0 m/s, with the exception of one participant who walked at 0.8 m/s due to perceived instability while wearing the experimental footwear. The first 5 minutes of walking were performed in the symmetrical rigid shoe condition, after which the maximum compliance condition was triggered on one side. After 10 minutes of exposure to asymmetry, we triggered the shoes to return to symmetrical rigid states. Of the 12 participants included in the study, 8 experienced a left side perturbation, and 4 experienced a right side perturbation. During this bout of walking, we recorded 3D ground reaction forces (GRF) from the treadmill and 3D marker coordinates, which we used to calculate weight bearing, propulsion, and spatio-temporal measures.

### D. Data analysis

We smoothed ground reaction force vectors and centers of pressure obtained from the instrumented treadmill with a fourth-order zero-lag Butterworth low-pass filter (30 Hz) using the filtfilt function in MATLAB (The Mathworks, Natick, MA). We identified heel strike and toe-off events for each limb as the moment in which the GRF magnitude for the corresponding force plate exceeded (heel strike) or fell beneath (toe-off) a threshold of 25 N. Gait events were manually verified for all strides, and strides in which the stance foot interacted with both force plates simultaneously were discarded.

Marker data were recorded at 100Hz with an eight-camera motion capture system (Miqus M3; Qualisys, Inc., Gothenburg, Sweden). Recorded marker data was labeled and processed in Qualisys software and then filtered with a fourth-order zero-lag Butterworth low-pass filter (6 Hz) using the filtfilt function in MATLAB to remove high frequency noise. We performed all experimental data processing with custom scripts using MATLAB.

### E. Dependent Measures

To quantify weight bearing, we calculated the first (weight acceptance) and second (push-off) vertical GRF (vGRF) peak, the minimum vGRF during mid-stance, and the time-integral of the vGRF profile during stance (vGRF impulse) per side, per stride.

To quantify propulsion, we calculated the peak propulsive GRF (pGRF) and pGRF impulse per side, per stride. We also calculated the peak and impulse of the braking GRF (bGRF) per side for all strides.

To quantify spatio-temporal parameters of gait, we calculated stance time per side as the elapsed time between heel strike and subsequent toe off events. We similarly calculated step time per side as the elapsed time between heel strike of the ipsilateral limb and subsequent heel strike of the contralateral limb. We calculated step length as the anteriorposterior distance between heel markers at the moment of heel strike of the corresponding side. Stride time and stride length are defined as the sum of the step time or step length of both sides for a given stride.

For all bilateral measures (i.e., excluding stride length and stride time), we calculated an asymmetry ratio to indicate the relative magnitude of a measure on the perturbed vs. unperturbed side [9], [20], [21]. This ratio is defined as:

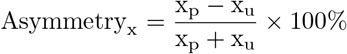

where *x*_p_ is the measure calculated for the side that was perturbed with compliance, and *x*_u_ is the measure for the side that remained unperturbed. Unless otherwise specified, all statistical analyses were performed on the asymmetry ratio of the corresponding measure.

### F. Statistical Analysis

We averaged all measures per participant in 10-stride windows to represent five experimental conditions: the final 10 strides of the baseline walking trial (“Baseline”), the first 10 strides of the perturbation, delayed by 4 strides to account for time required to achieve the compliance change (“Asym-early”), 10 strides at the end of the perturbation (“Asym-late”), 10 strides beginning 4 strides after the perturbation ended (“Post-early”), and the final 10 strides after 5 minutes of postperturbation walking (“Post-late”).

To determine whether asymmetric stiffness walking significantly influenced a given measure, we used a one-way repeated measures analysis of variance (ANOVA). We tested for sphericity (i.e., the assumption of equal variances of the differences between all combinations of conditions) with Mauchly’s test; if the assumption of sphericity was violated, we applied the Greenhouse-Geisser correction factor was to the degrees of freedom of the ANOVA. For measures with a significant effect of walking period condition, we conducted a post-hoc comparison using a two-tailed paired t-test between the Baseline and Post-early conditions to test our hypothesis that asymmetric walking would result in an asymmetry relative to baseline after the perturbation. We performed all statistical analyses using MATLAB (Statistics and Machine Learning Toolbox), and the significance level was set at *α* = 0.05 for all tests.

## III. RESULTS

### A. Weight bearing

We found a significant effect of condition in the pushoff vGRF peak (*F* (4, 44) = 5.47, *p* = .001) and in vGRF impulse (*F* (4, 44) = 12.95, *p <* .001), but not in weight acceptance (*F* (2.13, 23.50) = 1.25, *p >* .1) or mid-stance peaks (*F* (4, 44) = 1.95, *p >* .1). Relative to Baseline, participants had lower push-off vGRF peaks (–1.52% asymmetry, *t*(11) = ™3.18, *p* = .009) and vGRF impulse (–2.89% asymmetry, *t*(11) = ™4.13, *p* = .002) on the perturbed side than the unperturbed side during the Post-early window. An increase in negative asymmetry during Post-early supports our hypothesis that the perturbation would cause a shift in vertical ground reaction forces toward the unperturbed side.

**Fig. 2.**
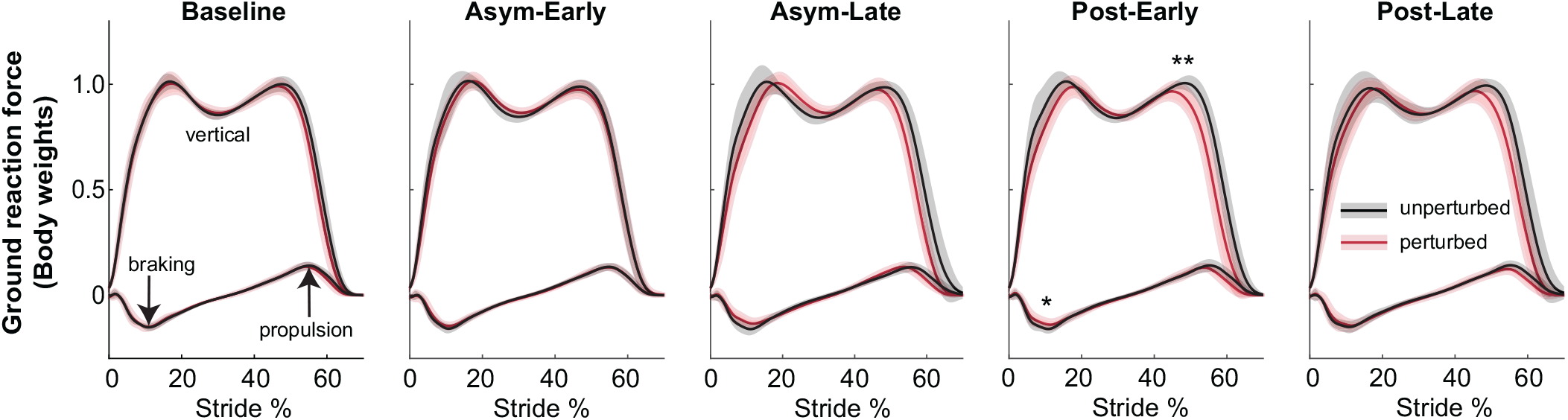
Ground reaction force (GRF) traces during Baseline, Asym-early, Asym-late, Post-early, and Post-late conditions. Per-stride transient measures are illustrated and labeled in the “Baseline” plots. Asterisks reflect significant difference to baseline via paired t-test: * = ***p <*** .**05**, ** = ***p <*** .**01**, *** = ***p <*** .**001**.

### B. Braking and propulsion

We found a significant effect of condition in peak braking (*F* (4, 44) = 4.63, *p* = .003), peak propulsion (*F* (2.65, 29.19) = 3.24, *p* = .041), and propulsion impulse (*F* (4, 44) = 3.93, *p* = .008), but not in braking impulse (*F* (4, 44) = 0.95, *p >* .1). Relative to Baseline, participants had lower peak braking force on the perturbed side than on the unperturbed side during the Post-early window (–7.04% asymmetry, *t*(11) = − 2.75, *p* = .019). This change in peak braking symmetry supports our hypothesis that the perturbation would elicit an adaptation toward increased dependence on the unperturbed leg for braking. Despite the significant effect of condition for peak propulsion and propulsion impulse, there was no significant difference in Post-early from Baseline for these measures (*p >* .1 for both). Counter to our hypothesis, we did not observe an adaptation that shifted propulsion forces from the perturbed to the unperturbed limb.

### C. Spatio-temporal

We found a significant effect of condition in step time (*F* (1.88, 20.67) = 4.16, *p* = .032), stance time (*F* (4, 44) = 10.39, *p <* .001), and step length (*F* (4, 44) = 5.79, *p* = .001), but not in stride time (*F* (4, 44) = 0.98, *p >* .1) or stride length (*F* (4, 44) = 1.81, *p >* 0.1). Relative to Baseline, participants walked with shorter stance time on the perturbed side than on the unperturbed side during the Post-early window (–1.03% asymmetry, *t*(11) = −2.71, *p* = .020). There was no significant difference between the Post-early and Baseline conditions for step time or step length (*p >* .1 for both). While we did not have a specific hypothesis about spatio-temporal adaptation, this result indicates a distinct difference between adaptation to the ACFT perturbation and split-belt treadmill walking, for which step length adaptation is a common observation [8], [22]–[24]. The significant observation of negative stance time asymmetry indicates a change in weight-bearing strategy that involves duration of contact with the ground, in addition to the GRF magnitude adaptations reported above.

**TABLE I.** Ground reaction force and spatiotemporal outcome measures.

|  | Baseline | Asym-early | Asym-late | Post-early | Post-late | Main effect p-value |
| --- | --- | --- | --- | --- | --- | --- |
| Peak weight acceptance vGRF (% Asym) | $-0.2 \pm 1.3$ | $-0.7 \pm 2.7$ | $-0.7 \pm 3.7$ | $-1.8 \pm 4.0$ | $-1.1 \pm 3.1$ | 0.306 |
| Peak push-off vGRF (% Asym) | $-0.3 \pm 1.3$ | $-0.4 \pm 1.7$ | $-0.8 \pm 1.6$ | $-1.8 \pm 2.0$ | $-1.3 \pm 2.3$ | **0.001** |
| Min. midstance vGRF (% Asym) | $0.2 \pm 1.3$ | $1.2 \pm 1.5$ | $1.5 \pm 1.8$ | $0.6 \pm 1.7$ | $0.7 \pm 1.5$ | 0.119 |
| vGRF impulse (% Asym) | $-1.1 \pm 1.6$ | $-0.5 \pm 2.1$ | $-3.3 \pm 2.9$ | $-4.0 \pm 3.2$ | $-3.3 \pm 2.8$ | **<0.001** |
| Peak braking (% Asym) | $0.1 \pm 6.1$ | $-2.7 \pm 6.7$ | $-8.9 \pm 12.1$ | $-7.0 \pm 10.8$ | $-1.9 \pm 8.7$ | **0.003** |
| Peak propulsion (% Asym) | $-1.9 \pm 5.7$ | $-0.9 \pm 6.2$ | $-2.0 \pm 10.0$ | $-3.6 \pm 10.6$ | $-7.2 \pm 7.9$ | **0.041** |
| Braking impulse (% Asym) | $-1.1 \pm 6.9$ | $-2.0 \pm 7.8$ | $-3.4 \pm 11.3$ | $-3.5 \pm 12.2$ | $0.0 \pm 10.3$ | 0.447 |
| Propulsion impulse (% Asym) | $-3.6 \pm 8.6$ | $-0.8 \pm 8.8$ | $-4.4 \pm 12.8$ | $-7.3 \pm 12.9$ | $-8.5 \pm 12.6$ | **0.008** |
| Step time (% Asym) | $-1.1 \pm 1.8$ | $0.1 \pm 1.9$ | $-1.8 \pm 3.2$ | $-1.9 \pm 3.4$ | $-2.3 \pm 2.6$ | **0.032** |
| Stance time (% Asym) | $-0.7 \pm 1.1$ | $0.1 \pm 1.5$ | $-1.6 \pm 1.7$ | $-1.7 \pm 2.1$ | $-1.6 \pm 1.5$ | **<0.001** |
| Step length (% Asym) | $0.0 \pm 2.4$ | $-0.9 \pm 3.7$ | $-0.3 \pm 4.5$ | $1.6 \pm 4.4$ | $2.3 \pm 3.8$ | **0.001** |
| Stride time (s) | $1.200 \pm 0.047$ | $1.194 \pm 0.047$ | $1.210 \pm 0.064$ | $1.201 \pm 0.056$ | $1.211 \pm 0.063$ | 0.430 |
| Stride length (m) | $1.022 \pm 0.093$ | $1.006 \pm 0.095$ | $1.024 \pm 0.113$ | $1.027 \pm 0.105$ | $1.028 \pm 0.116$ | 0.144 |

## IV. DISCUSSION

The goal of this study was to test whether prolonged perturbations to foot-ground compliance as delivered by active footwear would elicit signatures of neural adaptation in weight-bearing. Overall, participants exhibited significant weight-bearing aftereffects following exposure to asymmetric shoe compliance during walking, indicating that the perturbation elicited neural adaptation as expected. In support of our hypothesis, we observed a shift in weight-bearing measures away from the perturbed-side limb. Counter to our hypothesis, we did not observe a significant aftereffect in propulsion. Importantly, these results indicate that the dissipative nature of the active footwear plays a significant role in how observed gait behavior adapts to changes in stiffness.

### A. Weight-bearing adaptations

We observed aftereffects in peak vGRF during push-off and vGRF impulse asymmetry, suggesting that the shoe compliance perturbation elicited neuromotor adaptations to weight bearing. Both push-off vGRF and vGRF impulse shifted toward the unperturbed limb post-perturbation, indicating that participants adapted by relying more on the unperturbed limb to maintain pendular mechanics at the step-to-step transition and perform body support during stance, respectively. Both of these mechanical functions of the stance limb are deficient on the paretic side in hemiparetic stroke gait, making this a beneficial adaptation should it generalize to impaired neuromotor control.

As predicted by neuromuscular simulations [19], a dissipative compliance perturbation elicited a shift in weightbearing forces away from the perturbed limb. This avoidant strategy is somewhat intuitive; by making the foot-ground interaction on one side less stable and energetically inefficient, the perturbation effectively impairs the limb it is applied to. Anecdotally, some participants reported a perceived need to walk more carefully during the perturbation due to the loss of mediolateral stability on the perturbed side.

While we did not formally test for aftereffect duration, weight bearing asymmetry appears to not wash out immediately after the perturbation, with some vGRF asymmetry still visible after 5 minutes of post-perturbation walking (Fig. **??**). This is promising from a gait training perspective, although it is difficult to decouple lingering effects due to neuromotor adaptation from potential fatiguing effects of the perturbation. Future work will include formal analysis of aftereffect duration per measure.

### B. Braking and propulsion adaptations

We observed significant main effects in braking and propulsion measures, but the only significant aftereffect was in peak braking. As with the vGRF measures, peak braking shifted toward the unperturbed limb. While it appears that participants make adaptations to the relative role of each limb during push-off, the change does not translate to changes in propulsion symmetry. The presence of a significant aftereffect in peak braking but not in weight acceptance vGRF and an aftereffect in push-off vGRF but not in propulsion, suggests that changes occur during both both weight acceptance and push-off gait events, but that they are dependent on limb angle (i.e., participants are adjusting to push-off vertically instead of horizontally). This same pattern was observed in experiments with asymmetric surface stiffness treadmill walking [15], where increases in plantarflexor muscle activity were also observed for the limb with increased vertical push-off force. While we did not measure muscle activity in this study, we can speculate that the same pattern could be occurring here.

### C. Spatio-temporal symmetry adaptations

We observed significant main effects in step time, stance time, and step length, but no significant aftereffects for any spatio-temporal measure. While the aftereffect in stance time did not reach statistical significance, its pattern closely mirrors that of vGRF impulse, which did show a significant aftereffect.

**Fig. 3.**
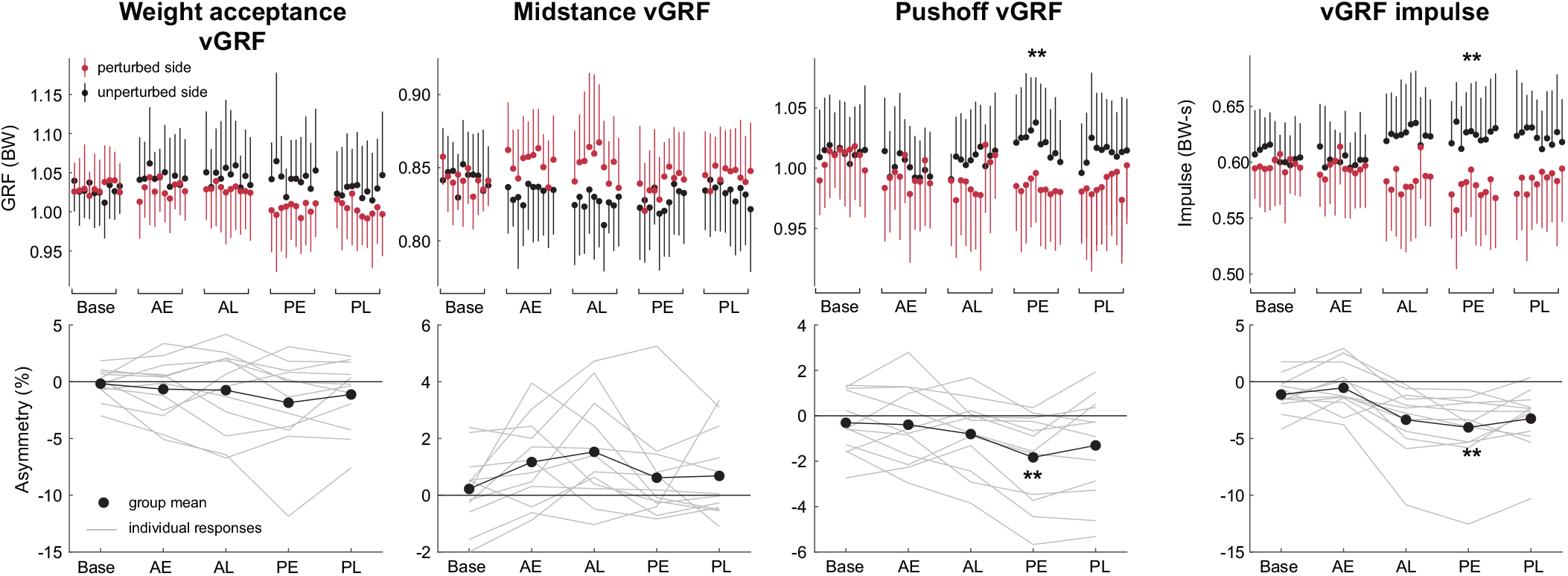
Group means of vertical GRF measures per condition (Base = Baseline, AE = Asym-early, AL = Asym-late, PE = Post-early, PL = Post-late) for (top) the perturbed and unperturbed limbs per stride and (bottom) asymmetry ratio. Error bars reflect one standard deviation. Asterisks reflect significant difference to baseline via 476 paired t-test: * = ***p <*** .**05**, ** = ***p <*** .**01**, *** = ***p <*** .**001**.

**Fig. 4.**
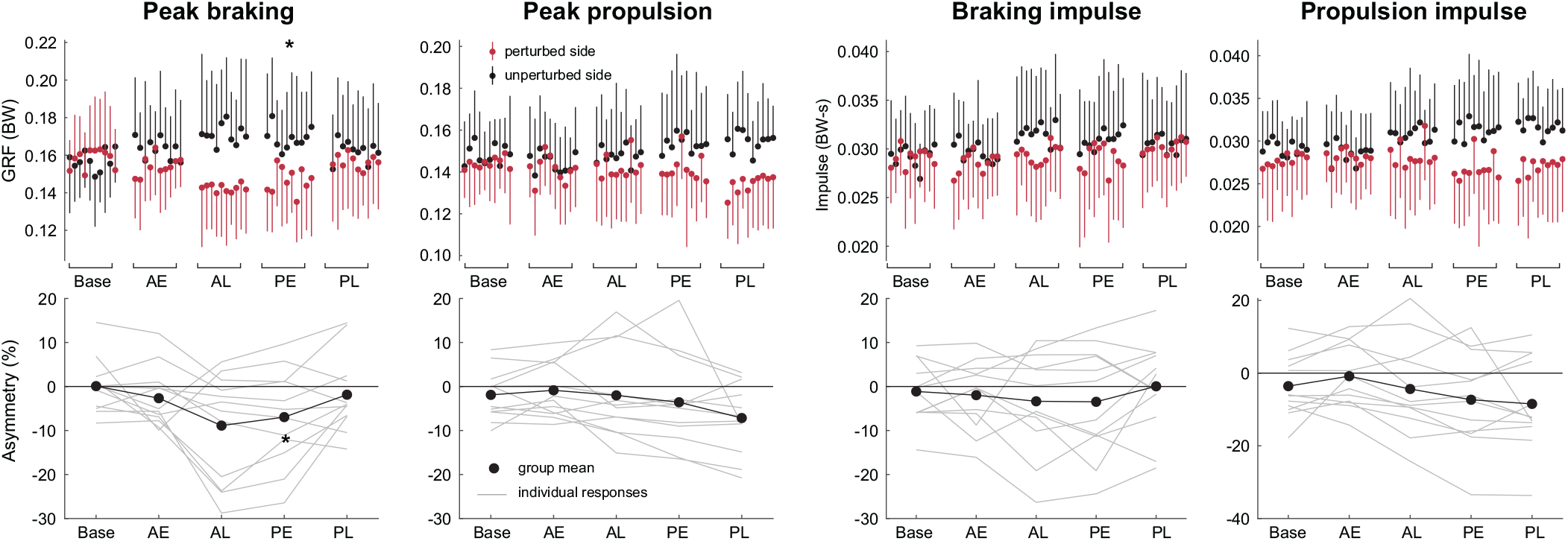
Group means of braking GRF and propulsive GRF measures per condition (Base = Baseline, AE = Asym-early, AL = Asym-late, PE = Post-early, PL = Post-late) for (top) the perturbed and unperturbed limbs per stride and (bottom) asymmetry ratio. Error bars reflect one standard deviation. Asterisks reflect significant difference to baseline via 476 paired t-test: * = ***p <*** .**05**, ** = ***p <*** .**01**, *** = ***p <*** .**001**.

This suggests that a true aftereffect in stance time may exist but is not detectable within the statistical power of the current study. Regardless, this is a distinct difference from splitbelt treadmill adaptation, in which there are typically clear spatio-temporal symmetry aftereffects but unclear aftereffects in ground reaction forces []. The small overlap in the postperturbation effects in split-belt treadmill adaptation and asymmetric ground compliance adaptation may in fact be beneficial, as it may allow the combination of these two perturbations with minimal unwanted interactions.

### D. Comparison to asymmetric stiffness treadmill adaptation

Participants showed similar patterns in their response to the footwear-applied perturbation compared to a treadmill-applied perturbation. As previously stated, both perturbations elicited aftereffects in push-off vGRF but not weight acceptance vGRF, and in braking vGRF but not propulsion vGRF. Critically, however, the direction of the elicited asymmetry was reversed for the footwear-applied perturbation; participants have previously been shown to shift their ground reaction forces toward the perturbed side after walking on an asymmetric stiffness treadmill [15]. One explanation for this difference is in the difference in mechanical dynamics between the two device types. The spring response of existing asymmetric stiffness treadmills is in large part driven by their inertia, which causes an underdamped response with a relatively low natural frequency [12], [13]. The compliant footwear, on the other hand, is highly dissipative and low in inertia, resulting in an overdamped response [18]. In support of this explanation, musculoskeletal simulations of walking on asymmetric surface dynamics predict larger peak vGRF on the perturbed side for low-damping perturbations and on the unperturbed side for high-damping perturbations [19]. These simulations minimized a measure for overall muscle effort, which also results in reproducing features of adapted split belt treadmill walking [25], [26]. It is possible that increasing the dissipative element of the perturbation shapes adaptation toward an avoidant strategy as opposed to one that exploits the perturbed dynamics.

In addition to reversing the direction of the aftereffects, the footwear perturbation elicited aftereffects in vGRF impulse symmetry which were not observed in a treadmill perturbation [15]. This measure represents the relative contribution of each limb to body support during gait, therefore eliciting a change in vGRF impulse symmetry may be important for future clinical applications. The difference of response in vGRF impulse between perturbation types may also be explained by the difference in energy conservation between the two types of compliant dynamics, but additional research into the compliance dynamics parameter space is required to rigorously explain this difference.

Finally, there is an apparent difference in the speed of the adaptation response to the footwear relative to adaptation to treadmill-based asymmetry perturbations. Asymmetric treadmill stiffness and belt speed both typically induce a large initial asymmetry in ground reaction forces and step length, respectively [15], [24]. While we did not formally evaluate adaptation or de-adaptation time, there is a distinct lack of this large initial response to a stiffness perturbation applied by the footwear. It is possible that due to the limited overall deflection magnitude and stiff low-loading response of the footwear [18], the perturbation was not destabilizing enough to require the fast time-scale adaptation response associated with maintaining upright stability, and was instead dominated by the slower timescale change associated with neuromotor adaptation [26], often theorized to be driven by energy optimization [20], [25]–[27]

### E. Study limitations and future work

Because the footwear perturbation requires a deformable interface to be attached to each foot, it is likely that there are some differences in walking mechanics relative to walking in normal footwear. However, these changes should affect both sides symmetrically, so we do not consider this to be a major factor in comparing symmetry changes between perturbation types. The wearable nature of the footwear perturbation allows for adaptation experiments to occur on a fully instrumented treadmill, albeit with a compliant interface between the force plate and the foot that may transiently distort the GRF measured by the force plate relative to the forces directly acting on the foot.

**Fig. 5.**
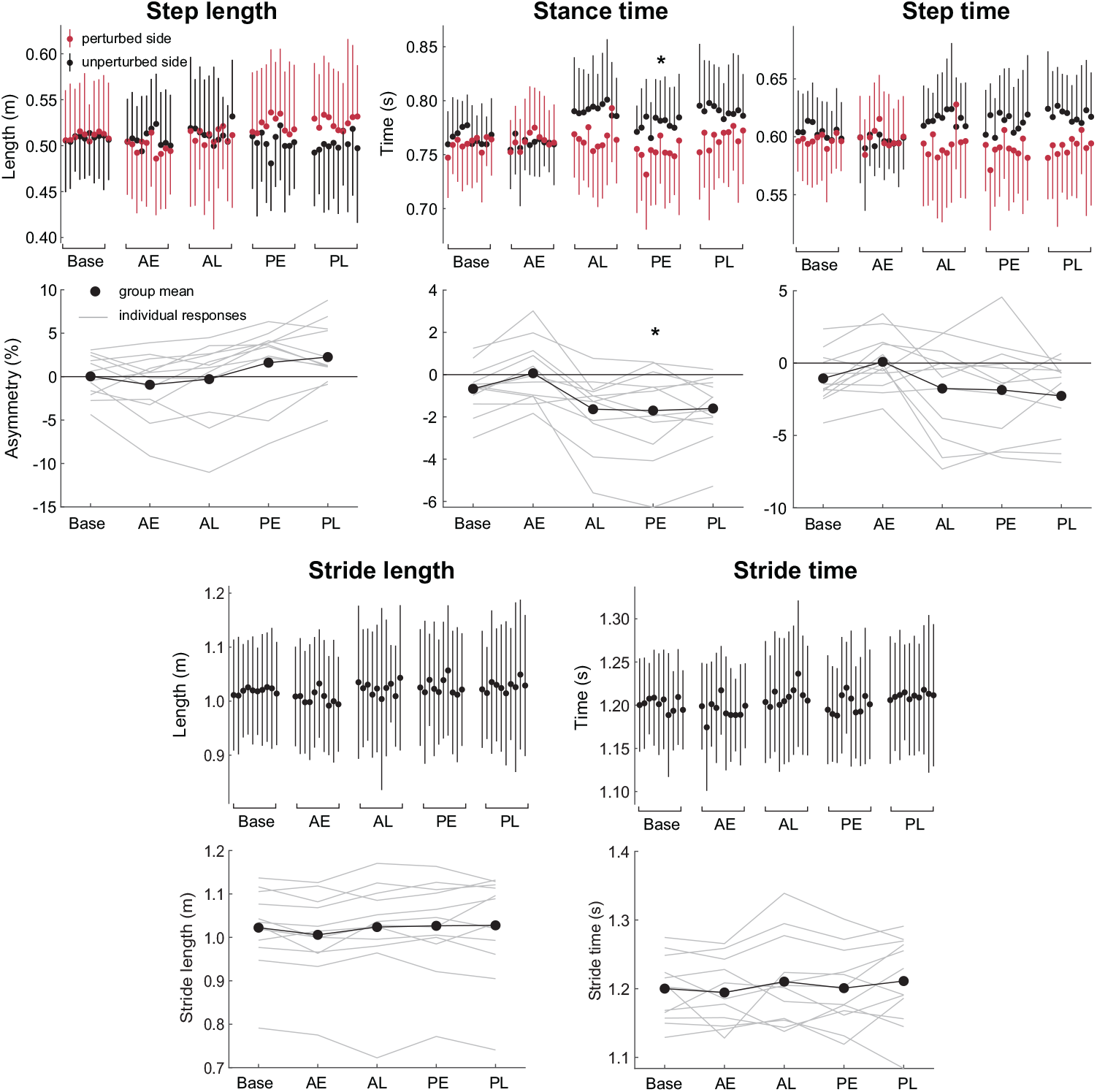
Group means of spatiotemporal measures per condition (Base = Baseline, AE = Asym-early, AL = Asym-late, PE = Post-early, PL = Post-late) (top) per stride and (bottom) asymmetry ratio, where relevant. Measures that comprise the sum of bilateral measures (stride length, stride time) are reported with their directly computed value rather than an asymmetry ratio. Error bars reflect one standard deviation. Asterisks reflect significant difference to baseline via 476 paired t-test: * = ***p <*** .**05**, ** = ***p <*** .**01**, *** = ***p <*** .**001**.

In this study, we focused on spatiotemporal and ground reaction force outcomes as they comprise our hypothesis and form a direct point of comparison with other asymmetric gait adaptation literature. The individual joint level and muscle level contributions to the observed behavior are also of interest to better explain the biomechanical outcomes and mechanisms for the adaptation. Future studies will quantify these additional measures.

To best characterize the adaptation and de-adaptation to an abrupt asymmetrical ground compliance perturbation, all of the experimental conditions involved the robotic footwear being worn. The effect of the perturbation on walking in normal shoes is also of interest, and future work will involve quantifying aftereffects with the robotic footwear removed. Furthermore, the findings from this study will form a comparison point for future work investigating overground adaptation.

## V. CONCLUSIONS

An asymmetric ground compliance perturbation applied via active footwear elicited changes to asymmetry in pushoff GRF and vertical GRF impulse, indicating a neuromotor adaptation to weight-bearing strategy during gait. We observed weight-bearing symmetry changes in the opposite direction to a similar intervention with an asymmetric surface stiffness treadmill, linked to differences in the mechanical dynamics between the two systems as predicted by optimal control simulations of walking using musculoskeletal models. Further study is needed to quantify asymmetric ground compliance adaptation overground using a footwear-based perturbation in comparison with adaptation confined to a treadmill.

## Acknowledgment

We would also like to thank Sarah Szemethy and Calder Robbins for their work making this experiment possible.

